# Single gene targeted nanopore sequencing enables simultaneous identification and antimicrobial resistance detection of sexually transmitted infections

**DOI:** 10.1101/2020.11.06.371310

**Authors:** Liqing Zhou, Andrea Lopez Rodas, Luz Marina Llangarí, Natalia Romero Sandoval, Philip Cooper, Syed Tariq Sadiq

## Abstract

**Objectives:** To develop a simple DNA sequencing test for simultaneous identification and antimicrobial resistance (AMR) detection of multiple sexually transmitted infections (STIs).

**Methods:** Real-time PCR (qPCR) was initially performed to identify *Neisseria gonorrhoeae* (NG), *Chlamydia trachomatis* (CT), *Mycoplasma genitalium* (MG) and *Trichomonas vaginalis* (TV) infections among a total of 200 vulvo-vaginal swab samples from female sex workers in Ecuador. qPCR positive samples plus qPCR negative controls for these STIs were subjected to single gene targeted PCR MinION-nanopore sequencing using the smartphone operated MinIT.

**Results:** Among 200 vulvo-vaginal swab samples 43 were qPCR positive for at least one of the STIs. Single gene targeted nanopore sequencing generally yielded higher pathogen specific read counts in qPCR positive samples than qPCR negative controls. Of the 26 CT, NG or MG infections identified by qPCR, 25 were clearly distinguishable from qPCR negative controls by read count. Discrimination of TV qPCR positives from qPCR negative controls was poorer as many had low pathogen loads (qPCR cycle threshold >35) which produced few specific reads. Real-time AMR profiling revealed that 3/3 NG samples identified had *gyrA* mutations associated with fluoroquinolone resistance, while none of the MG samples possessed *23S rRNA* gene mutations contributing to macrolide resistance.

**Conclusions:** Single gene targeted nanopore sequencing for diagnosing and simultaneously identifying key antimicrobial resistance markers for four common genital STIs shows promise. Further work to optimise accuracy, reduce costs and improve speed may allow sustainable approaches for managing STIs and emerging AMR in resource poor and laboratory limited settings.

## Introduction

Sexually transmitted infections (STIs) remain a major public health problem worldwide, with an estimated 357 million new cases of *Chlamydia trachomatis* (CT), *Neisseria gonorrhoeae* (NG), *Treponema pallidum* (TP) and *Trichomonas vaginalis* (TV) per year [1], all which commonly cause a genital discharge syndrome in men and women. High rates of the STI, *Mycoplasma genitalium* (MG), have also been reported worldwide [2] and is associated with genital discharge syndrome in men and reproductive sequelae in women [3]. All these STIs are generally curable with existing, effective single-dose antibiotic regimens but, if left undiagnosed and/or untreated, can result in serious long-term reproductive health sequelae, particularly for women. Antimicrobial resistance (AMR) among some STIs to multiple classes of antibiotics has spread rapidly in recent years. For NG, loss of extended spectrum cephalosporins as first line empirical treatment is a major concern [4] as circulating multi - and extensive-drug resistance clones have been detected internationally [5]. For MG, macrolide resistance is now widely but not universally reported with increasing rates of resistance to fluoroquinolones also detected [6]. These developments have made treatment, and particularly empirical treatment challenging.

New World Health Organization (WHO) guidelines reinforce the need to treat these STIs with the right antibiotic, at the right dose, and the right time to reduce spread and improve sexual and reproductive health. For both NG and MG, accurate and rapid diagnostics which also predict antibiotic susceptibility are likely to be needed to achieve this. Laboratory based nucleic-acid amplification tests (NAAT) for detection of NG, CT, TV and MG are the current gold standard for detection, providing high sensitivity and specificity. NAAT are widely used in high-income countries but are often unavailable in resource-poor settings. Culture-based antimicrobial susceptibility testing (AST) for NG remains gold standard for phenotypically predicting AMR and commonly takes usually two to five days to obtain a result in UK microbiology laboratories (personal communication, Sadiq). This in practical terms may be too late to initiate targeted antibiotic therapy, particularly for hard to reach vulnerable populations. MG is difficult to cultivate and requires cell culture, not usually feasible in most clinical settings.

Advances in understanding AMR in NG, MG and TV have allowed for development of NAAT-based AMR detection. For example, absence of *gyrA* mutations in NG accurately predicts fluoroquinolone susceptibility [7,8], presence of *23S rRNA* mutations in MG is associated with failure of treatment with azithromycin [9–11], and *ntr6* mutations in TV may have diagnostic value for metronidazole resistance [12]. Use of NAAT-based AMR tests, however, may have limitations due to continually changing mutations and novel mechanisms of resistance evolving under ongoing treatment selection pressures. Whole genome sequencing (WGS) using high throughput sequencing platforms may address this challenge to some degree and give added value in identifying phylogenetic relationships in identified infections [13,14]. However, for diagnostic purposes WGS itself can be constrained by sample preparation, cost etc., making it unsuitable for near patient applications.

Oxford Nanopore Technologies’ (ONT) portable MinION DNA sequencer, together with its recent MinIT hand-held processor, may offer advantages as an accurate diagnostic in resource-limited settings. Herein we report early phase study among female sex workers (FSWs) in Ecuador, of using the MinION, controlled by a smartphone-operated MinIT to detect the four common STIs and profile AMR to ciprofloxacin in NG, azithromycin in MG and metronidazole in TV utilising a single gene targeted approach.

## Materials and Methods

### Ethics statement

Written and verbal informed consent was obtained from FSW participants all of whom were 18 years or older and none of whom received any compensation for their participation. The study was conducted according to the Declaration of Helsinki and approved by the Ethical Committee of Universidad Internacional del Ecuador (02-02-17).

### Clinical sample collection and DNA preparation

Vulvo-vaginal swab samples were collected by clinicians using Xpert^®^ CT/NG Patient-Collected Vaginal Swab Specimen Collection Kit (Cepheid). DNA from swab samples was prepared using PureLink Genomic DNA Mini Kit (Thermo Fisher Scientific) according to the manufacturer’s instructions and quantified using a NanoDrop Spectrometer.

### STI identification of swab samples by qPCR

Swab samples were initially screened by qPCR to identify NG, TV, MG and CT infections, with gene targets and primers shown in Table 1. qPCR was performed using Applied Biosystems 7500 Fast Real-Time PCR System in a volume of 10 μl containing 5 μl of TaqMan™ Fast Universal PCR Master Mix, 1 μl of 10x Exogenous Internal Positive Control (IPC) Mix, 0.2 μl of 50x IPC DNA, 250 nM each of the primers, 100 nM each of the probes and 50 ng of template DNA. Cycling parameters were 95 °C for 10 min, followed by 40 cycles of 95°C for 15 sec, 60°C for 1 min.

**Table 1.**
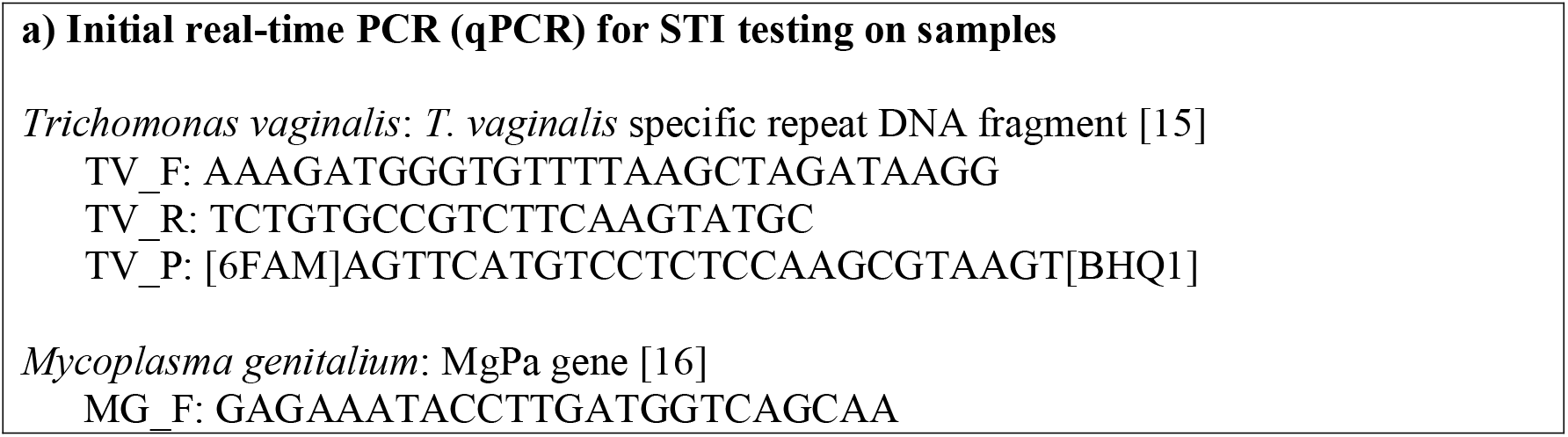
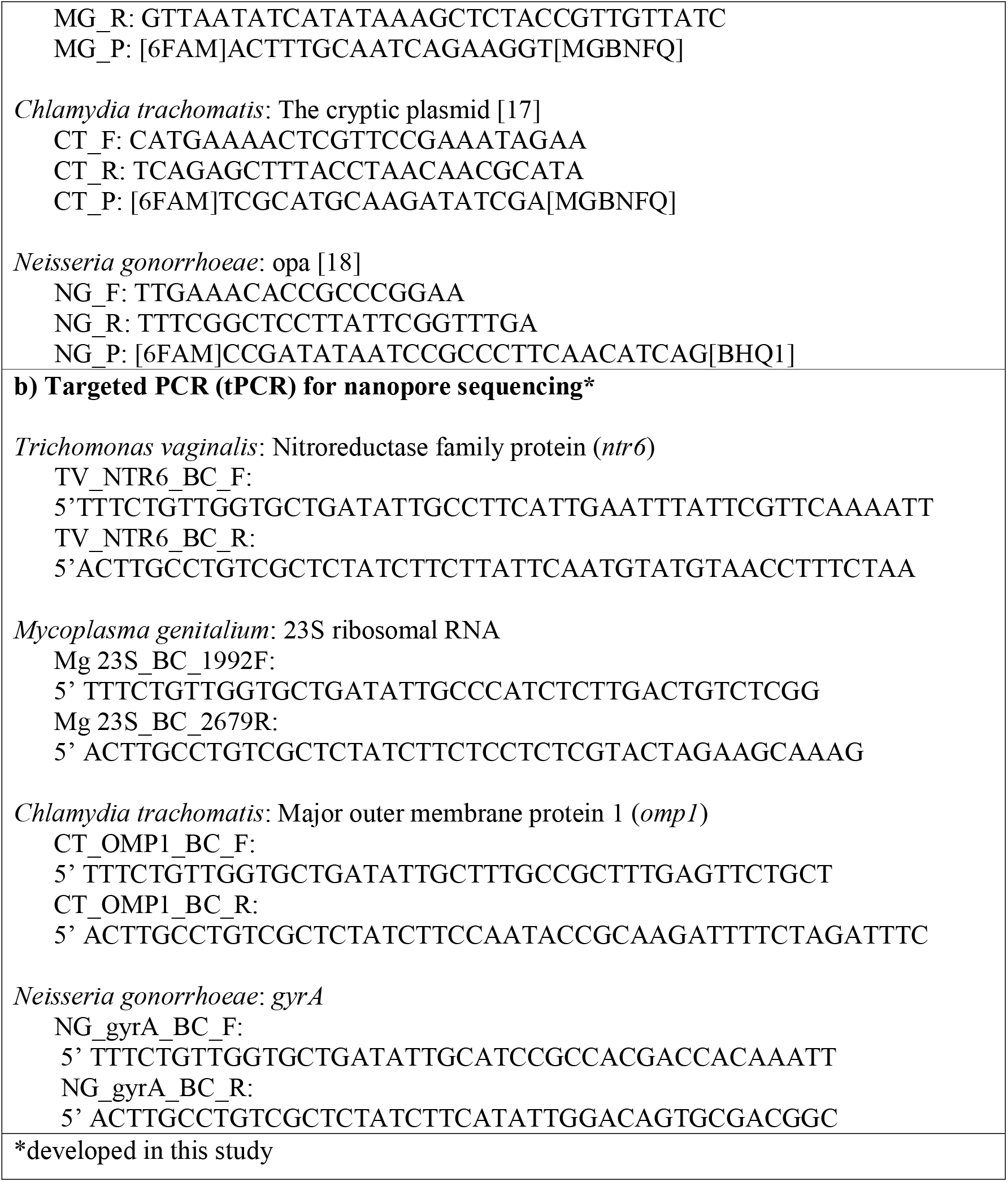
Gene targets, primers and probes used in this study

### Targeted PCR (tPCR) amplification and barcoding

Both qPCR positive and negative samples for NG, MG, TV and CT plus an NG positive control were amplified by tPCR, targeting genes *gyrA* (NG), *23S rRNA* (MG), *ntr6* (TV) and *omp1* (CT) with the primers listed in Table 1. tPCR reactions for each target were performed in a 50 μl volume consisting of 25 μl LongAmp^®^ Taq 2X Master Mix (NEB), 200 nM each of tPCR primers and 100 ng DNA template as follows: 95 °C for 3 min followed by 35 cycles of 95 °C for 30 sec, 58 °C for 30 sec, 65 °C for 1 min 30 sec, and one cycle of 65 °C for 5 min. tPCR amplicons were purified and subjected to a second PCR for barcoding using PCR Barcoding Expansion 1-96 (EXP-PBC096, ONT) according to the manufacturer’s instructions.

### Nanopore sequencing and data analysis

A DNA sequencing library was constructed from a pool of barcoded tPCR amplicons of different clinical samples using Ligation Sequencing Kit (SQK-LSK108) and sequenced using FLO-MIN106 R9 Version Flow Cell MK I Spot-ON on portable DNA sequencer MinION MK I which was controlled by a smartphone operated MinIT (ONT) according to the manufacturer’s instructions. Sequencing data were uploaded onto Metrichor Epi2ME (ONT) and analyzed using the workflow Fastq Antimicrobial Resistance r3.3.2 which included three components: QC and Barcoding [rev. 3.10.4], WIMP [rev. 3.4.0] and ARMA CARD [rev. 1.1.6]. QC and Barcoding component contains quality score (qscore) and barcode filter, cutting off reads with a qscore below the threshold (min qscore:7) which will not be uploaded for analysis, and demultiplexing barcodes if the sequenced samples are barcoded. WIMP (what’s In My Pot) allows for species identification by classifying read sequences against the standard Centrifuge database (including RefSeq complete genomes for bacteria). ARMA (antimicrobial resistance mapping application) performs antimicrobial resistance identification by aligning input reads with minmap2 against all reference sequences available in the CARD database (CARD version 1.1.3). TV, an anaerobic, flagellated protozoan parasite, was not included in RefSeq for bacteria and therefore a TV G3 nitroreductase family protein (TVAG_354010, ntr6) reference was uploaded using Fasta Reference Upload r3.2.2 and TV identification was performed using Fastq Custom Alignment r3.2.2 against the *ntr6* reference sequence. Read counts were log-transformed and compared between qPCR positive and negative samples by t-test.

### BLAST search

DNA sequences of NG, MG and TV reads were extracted from read FASTQ files and BLAST-aligned against the reference sequences: *Neisseria gonorrhoeae* FA 1090 *gyrA* reference (NC_002946.2:c621189-618439), *Mycoplasma genitalium* strain G-37 *23S rRNA* gene (NR_077054.1) and *Trichomonas vaginalis* G3 nitroreductase family protein reference (TVAG_354010) respectively. Part of the aligned sequences flanking AMR-associated mutations were used to construct a consensus sequence using MultAlin [19]. AMR-associated mutations were confirmed by manually comparing the consensus sequence against the reference.

## Results

### STI identification of swab samples by qPCR

A total of 200 vulvo-vaginal swab samples from FSWs were initially screened by qPCR, and 43 were found positive, including 37 single infections (11 MG, 19 TV and 7 CT) and six co-infections (two CT/TV and one each of CT/NG, NG/TV, CT/NG/TV and CT/TV/MG), as shown in Table S1 (Supplementary data).

### STI identification and AMR detection by single gene tPCR nanopore sequencing

All 43 qPCR positives plus 12 negatives (3 for each pathogen) and an NG positive control were subjected to single gene tPCR nanopore sequencing in one barcoded sequencing library. A sequencing run of 10 hours produced 1,499,872 reads of which 1,127,703 (76%) reads passed QC, with total yield of 1.2 gigabases, average qscore 8.34 (Figure S1, Supplementary data) and average sequence length 816 bases. To identify MG, NG and CT, all reads (1,127,703) which passed QC were analyzed by WIMP, resulting in 810,683 (72%) classified reads of which 43,491 (5.4%) were non-barcoded. For TV identification, Fastq Custom Alignment workflow produced 148,458/1,127,703 (13%) read alignments with TV *ntr6* sequence of which 66,500 (5.9%) were non-barcoded.

Targeted amplicon sequencing from clinical samples, as expected, still non-specifically produced some reads from other microbial and/or host DNA in the sample. However, for STI pathogens, qPCR positive samples generally yielded higher specific read counts than qPCR negative controls, all of the latter of which had fewer than 15 reads (Figure 1). Mean log_10_ read counts were higher in qPCR positive NG, MG and CT samples compared to negative by t-test (Table S2, Supplementary data) but not significantly for TV as 14/24 TV read counts in TV qPCR positive samples were very low (<20 reads). NG, MG and CT read counts clearly separated qPCR positive and negative samples except for one MG qPCR positive sample that had an absolute read count of 1 (Figure 1). The samples having <20 reads generally had a low load of pathogen, as evidenced by qPCR cycle threshold (Ct) being >35 (Table S1, Supplementary data).

**Figure 1.**
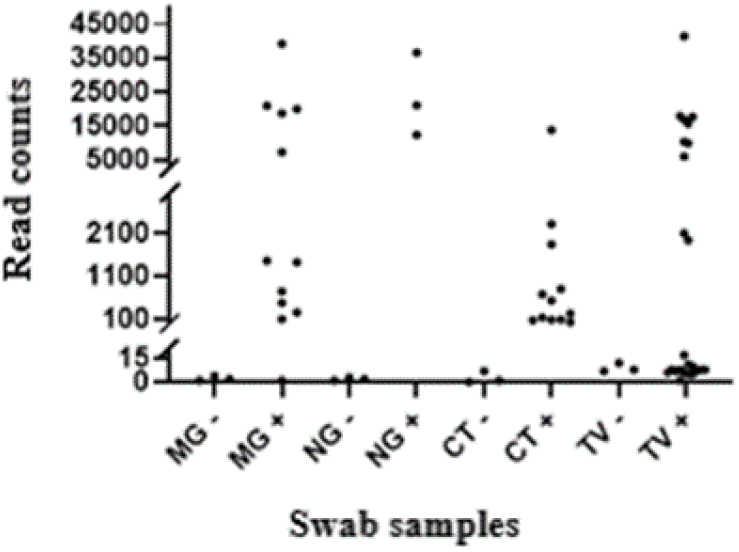
Pathogen read counts in the qPCR positive and negative STI samples analyzed by single gene targeted PCR nanopore sequencing

The ARMA workflow assigned 86,430/810,683 (11%) classified reads to antibiotic resistance genes in the CARD database, >99.99% of which were identified to be fluoroquinolone resistant *gyrA* gene in NG, using protein variant model. This analysis indicated that 3/3 NG clinical strains identified were fluoroquinolone-resistant, while none of the MG strains were resistant to macrolide antibiotics. The NG positive control, phenotypically susceptible to fluoroquinolones, and genotypically wild-type by BLAST (see below), was mis-identified to be fluoroquinolone-resistant. TV antimicrobial resistance was not profiled due to the absence of TV *ntr6* entries in the databases.

### AMR confirmation by manual BLAST search

A manual BLAST search was performed to ascertain the accuracy of the pathogen AMR identified by the ARMA workflow. The BLAST confirmed, as shown in Table 2, that all (3/3) of NG had fluoroquinolone-resistant mutations, 2/10 (20%) of TV had mutations related to metronidazole resistance, and none of MG had macrolide resistance-associated mutations. It also showed that the gyrA of NG positive control did not possess fluoroquinolone-resistant mutations.

**Table 2.**
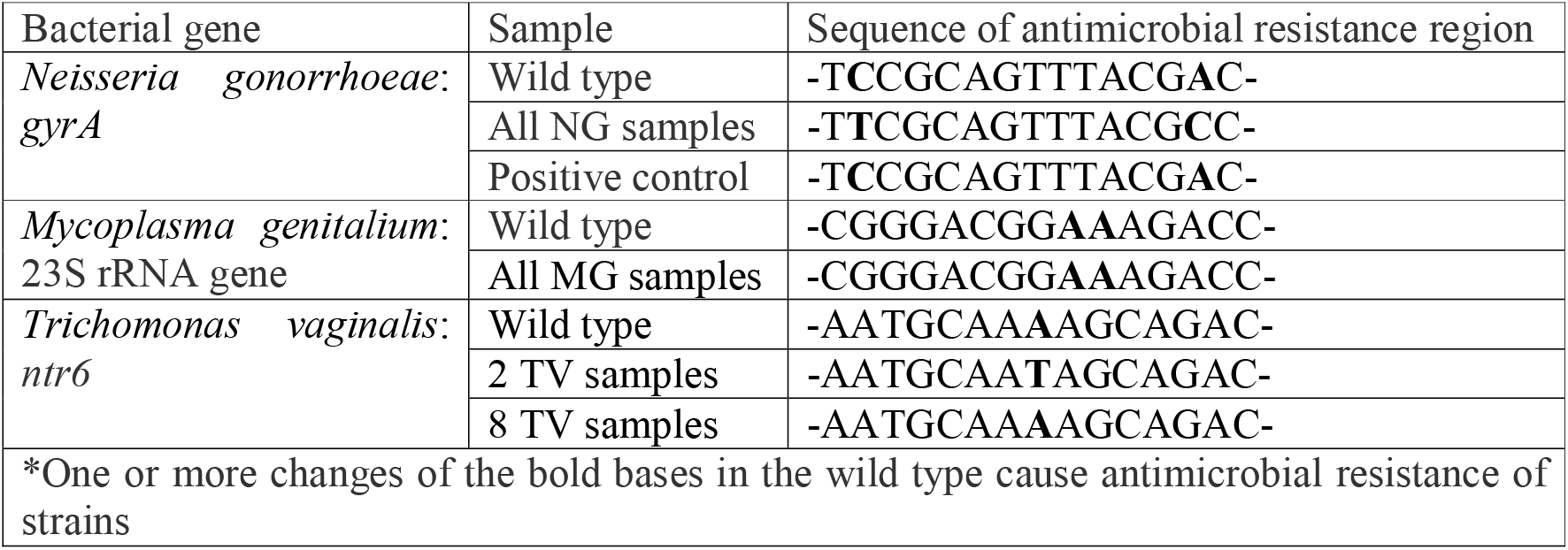
BLAST confirmation of antimicrobial resistance associated mutations identified by single gene targeted PCR nanopore sequencing*

## Discussion

We demonstrated the potential to test four common genital STIs and simultaneously detect AMR in three of them among vulnerable FSWs in Quito, Ecuador using the portable MinION DNA sequencer, controlled by smartphone, through a single gene targeted approach.

Single gene approaches for detection of infection, such as 16S rRNA sequencing have been increasingly described [20,21], particularly with the advent of long read sequencing such as PacBio and ONT [22], which has greater likelihood of taxonomically identifying organisms at species level. We intentionally evaluated the accuracy of sequencing of a single AMR gene, as opposed to a two-stage gene approach [23], to both diagnose infection and predict AMR simultaneously in order to evaluate potential for use in resource poor field settings where simplicity and cost become increasingly important factors to consider.

Our approach appeared to have some advantages over NAAT. We demonstrated initially that genes responsible for AMR have potential to be as useful as 16S rRNA genes for diagnosis and thus may serve as a ‘two-in-one’ target. We chose for both pathogen and AMR detection the NG *gyrA*, MG *23S rRNA* and TV *ntr6*, all gene mutations in which have been shown to be predictive of AMR [7–12,24,25], and CT *omp-1* for CT detection [26]. As sequencing allows for identification of evolving mutations in the same genes without changing the test, our approach will therefore have value as it is adapted to other gene targets undergoing continuous evolution under selection pressure such as penA in which changes may give rise to penicillin and extended spectrum cephalosporin resistance [27]. The approach can also allow for sequencing multiple genes simultaneously alongside vigilance over any changing nature of phenotypic to genotypic association for antibiotics where AMR involves multiple mechanisms.

Clinical STI samples contain a wide range of pathogen loads being drawn frequently from largely asymptomatic patients as in the case of the FSW cohort in this study. We found that except for TV, diagnosis by the single gene targeted nanopore sequencing appeared to be hardly limited by pathogen loads. In fact, our single gene approach correctly identified STI infections in the samples which had qPCR Ct<35 (higher pathogen loads). In the group with lower pathogen loads (Ct>35), tPCR amplification appeared not to enhance the absolute read counts of pathogen targets, possibly due to low tPCR efficiency caused by the presence of a large quantity of background DNA and non-optimal tPCR conditions. Further optimization of tPCR conditions may maximize the diagnostic sensitivity for this group of samples.

Regarding the Fastq Antimicrobial Resistance workflow used in this study, the component WIMP correctly identified each pathogen of interest, but we found the component ARMA CARD appeared to mis-profile the AMR of the NG control strain. This might be because the ARMA CARD used individual reads for matching AMR associated mutations, instead of a consensus sequence, supported by the result obtained with our manual BLAST search using the consensus sequence. Any further development of ARMA could help improve accuracy of AMR prediction.

This study had some limitations. Firstly, there was a small sample size, which impacted particularly on measures of specificity. Secondly no further confirmatory sequencing was performed to confirm the mutations present, largely due to the local regulations which prevented transporting material out of Ecuador. Thus, this study should be regarded as a proof of concept study. Future work is required on larger sample sets with composite reference standards in a formal diagnostic evaluation. Optimisation of amplification and target multiplexing will also be important.

Nanopore sequencing with MinION requires substantially lower infrastructure and startup cost compared to other high throughput sequencing platforms. Although arguably consumables are not optimally priced for resource poor settings, the promise of combining automated library preparations and use of disposable flow-cells has potential to use nanopore sequencing for field diagnostics. We recently demonstrated that even in developed settings implementation of rapid ciprofloxacin NAAT resistance tests for NG still requires net investment [28]. More recently ONT has developed a tablet version, MinION™ Mk1C which enables full sequencing and analysis to be performed in the lab and field without need for internet connection. This development could reduce cost and improve speed further to use nanopore sequencing as a diagnostic tool. However, this will be formally evaluated in a large-scale clinical study.

In conclusion, this study demonstrated that single gene targeted nanopore sequencing for diagnosing and simultaneously identifying key antimicrobial resistance markers for four common STIs shows promise. Further work to optimise accuracy, reduce costs and improve speed may allow sustainable approaches for managing STIs and emerging AMR in resource poor and laboratory limited settings.

## Contributions

STS and LZ conceived the study. NRS, PC and STS conceived the wider study for FSWs and developed protocols for recruitment of patients with LML and ALR. ALR, LZ and LML performed qPCR. LZ performed tPCR and nanopore sequencing. LZ and STS performed analysis. LZ and STS wrote the paper with contributions from all authors.

## Acknowledgements

The authors would like to thank the FSWs associations for supporting the development of this study.

## Funding

This work was supported by University Internacional del Ecuador (grant number UIDE-EDM04-2016-2017), the Wellcome Trust Institutional Strategic Support Fund (grant number 204809/Z/16/Z) awarded to St. George’s, University of London. The sponsors had no role in the study design, data collection, and analysis, or in the decision to prepare these data for publication.

## Transparency declarations

LZ and STS are inventors on patents in the field of STI diagnostics.

## Notes

### Summary of Updates

1. Supplemental file was updated; 2. The word qPCR was added to positive and negative samples to make it clear from the targeted PCR samples throughout the text.

