## Supplementary material for "Single gene targeted nanopore sequencing enables simultaneous identification and antimicrobial resistance detection of sexually transmitted infections": Table S1

**Table S1 Identification of swab samples with STIs by real-time PCR and Nanopore sequencing**

| **Sample ID** | **NG** | **CT** | **TV** | **MG** | **Ct value** | **Read counts** | | | | |
| --- | --- | --- | --- | --- | --- | --- | --- | --- | --- | --- |
|  |  |  |  |  |  | **Analysed** | **MG** | **CT** | **NG** | **TV** |
| 60 | - | - | - | + | 32.505 | 784 | 91 | 0 | 1 | 1 |
| 166 | - | - | - | + | 30.436 | 25188 | 19968 | 0 | 2 | 9 |
| 85 | - | - | - | + | 35.403 | 1931 | 477 | 3 | 1 | 4 |
| 31 | - | - | - | + | 34.976 | 7988 | 1416 | 2 | 0 | 4 |
| 230 | - | - | - | + | 28.800 | 23263 | 18676 | 2 | 0 | 5 |
| 103 | - | - | - | + | 33.375 | 18383 | 7262 | 0 | 3 | 5 |
| 13 | - | - | - | + | 32.980 | 26039 | 20902 | 0 | 0 | 9 |
| 109 | - | - | - | + | 29.870 | 39248 | 39248 | 0 | 2 | 7 |
| 239 | - | - | - | + | 36.210 | 3133 | 250 | 0 | 1 | 5 |
| 69 | - | - | - | + | 35.810 | 3412 | 737 | 0 | 0 | 9 |
| 263 | - | - | - | + | 29.592 | 2446 | 1457 | 0 | 1 | 2 |
| 260 | - |  |  | + | 36.672 | 633 | 1 | 1 | 1 | 4 |
| 51 | - | - | - | - |  | 2952 | 4 | 1 | 2 | 1 |
| 75 | - | - | - | - |  | 589 | 2 | 2 | 0 | 4 |
| 77 | - | - | - | - |  | 252 | 1 | 0 | 2 | 0 |
| 24 | - | - | + | - | 20.607 | 20815 | 2 | 0 | 3 | 10368 |
| 58 | - | - | + | - | 25.250 | 23714 | 2 | 0 | 3 | 17712 |
| 235 | - | - | + | - | 36.170 | 29942 | 4 | 3 | 1 | 5 |
| 256 | - | - | + | - | 39.995 | 25061 | 2 | 1 | 2 | 8 |
| 146 | - | - | + | - | 39.695 | 68032 | 4 | 0 | 3 | 8 |
| 144 | - | - | + | - | 24.409 | 22359 | 0 | 2 | 1 | 6026 |
| 145 | - | - | + | - | 38.037 | 32349 | 4 | 0 | 2 | 10 |
| 261 | - | - | + | - | 38.596 | 37283 | 6 | 0 | 2 | 8 |
| 90 | - | - | + | - | 23.581 | 17230 | 4 | 1 | 2 | 9824 |
| 106 | - | - | + | - | 39.839 | 18291 | 2 | 2 | 0 | 8 |
| 92 | - | - | + | - | 40.00 | 19009 | 0 | 0 | 1 | 1 |
| 72 | - | - | + | - | 36.897 | 20163 | 2 | 0 | 2 | 11 |
| 140 | - | - | + | - | 35.249 | 22397 | 6 | 0 | 1 | 6 |
| 253 | - | - | + | - | 37.778 | 30217 | 5 | 0 | 1 | 5 |
| 41 | - | - | + | - | 38.867 | 10297 | 11 | 2 | 2 | 7 |
| 124 | - | - | + | - | 27.540 | 23714 | 7 | 1 | 2 | 17882 |
| 71 | - | - | + | - | 27.565 | 4949 | 5 | 1 | 2 | 2101 |
| 258 | - | - | + | - | 38.636 | 14925 | 5 | 10 |  | 17 |
| 105 | - | - | + | - | 30.327 | 9029 | 8 | 2 | 4 | 1937 |
| 221 | - |  | + | - | 38.171 | 2513 | 2 | 1 | 1 | 6 |
| 89 | - |  | + | - | 35.079 | 30401 | 5 | 0 | 2 | 7 |
| 91 |  | - | + | - | 21.186 | 29511 | 2 | 1 | 1 | 16907 |
| 39 |  |  | + | - | 21.381 | 41735 | 0 | 0 | 1 | 41486 |
| 260 | - |  | + |  | 27.556 | 35209 | 0 | 1 | 0 | 15706 |
| 79 | - | - | - | - |  | 28262 | 1 | 0 | 1 | 12 |
| 84 | - | - | - | - |  | 6617 | 4 | 0 | 3 | 8 |
| 98 | - | - | - | - |  | 21575 | 4 | 0 | 3 | 7 |
| 260 | - | + |  |  | 35.905 | 17328 | 6 | 82 | 5 | 3 |
| 94 | - | + | - | - | 29.812 | 12745 | 3 | 530 | 3 | 4 |
| 259 | - | + | - | - | 30.331 | 8309 | 6 | 672 | 2 | 8 |
| 242 | - | + | - | - | 31.946 | 8443 | 3 | 1840 | 1 | 10 |
| 236 | - | + | - | - | 35.936 | 10335 | 8 | 65 | 3 | 7 |
| 157 | - | + | - | - | 30.126 | 10277 | 4 | 25 | 1 | 9 |
| 114 | - | + | - | - | 30.464 | 5086 | 1 | 2317 | 1 | 6 |
| 6 | - | + | - | - | 31.232 | 7582 | 24 | 127 | 1 | 18 |
| 221 | - | + |  | - | 35.968 | 2978 | 3 | 72 | 2 | 4 |
| 89 | - | + |  | - | 29.932 | 6565 | 3 | 220 | 0 | 10 |
| 30 |  | + | - | - | 30.886 | 14878 | 1 | 13796 | 2 | 3 |
| 39 |  | + |  | - | 30.536 | 999 | 1 | 797 | 1 | 3 |
| 102 | - | - | - | - |  | 3095 | 1 | 7 | 0 | 5 |
| 107 | - | - | - | - |  | 4664 | 2 | 1 | 2 | 7 |
| 125 | - | - | - | - |  | 101 | 2 | 0 | 1 | 9 |
| 30 | + |  | - | - | 19.033 | 27210 | 0 | 0 | 21087 | 1 |
| 91 | + | - |  | - | 20.035 | 39223 | 0 | 1 | 36590 | 5 |
| 39 | + |  |  | - | 21.504 | 28317 | 0 | 3 | 12325 | 3 |
| 126 | - | - | - | - |  | 14121 | 1 | 0 | 2 | 2 |
| 189 | - | - | - | - |  | 9168 | 5 | 1 | 3 | 1 |
| 193 | - | - | - | - |  | 11334 | 2 | 0 | 1 | 3 |
| NG positive control |  |  |  |  |  | 7937 | 0 | 0 | 7659 | 1 |
