## Supplementary material for "Single gene targeted nanopore sequencing enables simultaneous identification and antimicrobial resistance detection of sexually transmitted infections": Table S2

**Table S2 Mean log(10) read counts in clinical swab samples analysed by single gene targeted PCR nanopore sequencing by t-test**

| Pathogen | Mean Log Pos | Mean Log Control | t | p |
| --- | --- | --- | --- | --- |
| CT | 2.56 | 0.40 | 4.46 | 0.0006 |
| MG | 3.16 | 0.49 | 3.61 | 0.003 |
| NG | 4.33 | 0.46 | 23.87 | <0.0001 |
| TV | 2.19 | 0.99 | 1.28 | 0.21 |
