## Supplementary figures and images for "Single gene targeted nanopore sequencing enables simultaneous identification and antimicrobial resistance detection of sexually transmitted infections"

### Figure S1

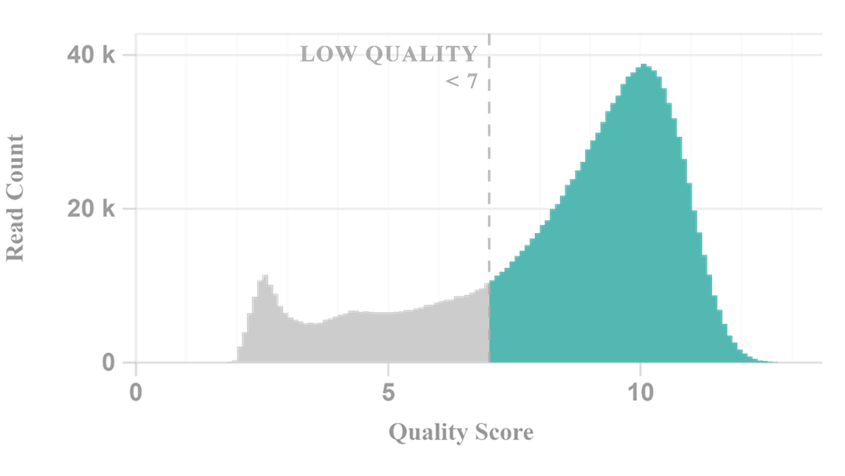

**Figure S1 Quality Scores of reads produced by single gene targeted PCR nanopore sequencing**
